# CpG Transformer for imputation of single-cell methylomes

**DOI:** 10.1101/2021.06.08.447547

**Authors:** Gaetan De Waele, Jim Clauwaert, Gerben Menschaert, Willem Waegeman

## Abstract

**Motivation:** The adoption of current single-cell DNA methylation sequencing protocols is hindered by incomplete coverage, outlining the need for effective imputation techniques. The task of imputing single-cell (methylation) data requires models to build an understanding of underlying biological processes.

**Results:** We adapt the transformer neural network architecture to operate on methylation matrices through combining axial attention with sliding window self-attention. The obtained CpG Transformer displays state-of-the-art performances on a wide range of scBS-seq and scRRBS-seq datasets. Furthermore, we demonstrate the interpretability of CpG Transformer and illustrate its rapid transfer learning properties, allowing practitioners to train models on new datasets with a limited computational and time budget.

**Availability and Implementation:** CpG Transformer is freely available at https://github.com/gdewael/cpg-transformer.

## 1 Introduction

DNA methylation is the addition of a methyl group to the DNA. The best-known type is CpG methylation, where the methyl group is added to the C-5 position of CG dinucleotides. Its association with a broad range of biological processes, such as gene expression regulation, is well-established [1]. CpG methylation is also known as a driving factor in developmental biology and carcinogenesis, motivating the need to study this phenomenon on the cellular level [2].

The last decade, several protocols that measure DNA methylation at single-cell resolution have been developed. These methods make use of bisulfite conversion of DNA followed by sequencing [3], both on genome-wide scale (scBS-seq) [4] and using reduced-representation protocols (scRRBS-seq) [5]. These methods have uncovered the heterogeneity and dynamics of epigenetic patterns between cells and have made it possible to describe epigenomic networks on an unprecedented scale and resolution [6].

Due to the smaller amount of genetic material available per cell, profiling single cells comes with certain challenges not encountered in bulk sequencing experiments. In practice, the genome-wide coverage of CpG sites per cell is low, ranging from 1% for high-throughput studies [7] to 20% for low-throughput ones [4]. Furthermore, profiled sites are covered by a smaller number of reads, resulting in noisy measurements of DNA methylation. Effective imputation and denoising techniques are therefore crucial in unlocking the full potential of single-cell methylome analyses.

Prediction of methylation states in tissue samples is a well-established problem in bioinformatics, often tackled by leveraging dependencies between CpG sites. For example, variational autoencoders have been successfully applied for dimensionality reduction of methylation data [8]. Other methods focus on imputation of single CpG sites in tissue samples using, among others, linear regression [9], Random Forests [10], autoencoders [11], gradient boosting [12] or mixture models [13]. In addition to using intrasample dependencies between neighboring CpG sites, some of these methods adopt the idea of leveraging information from multiple (tissue) samples for prediction [12, 13].

Most recent work on single-cell DNA methylation imputation has built upon this idea of leveraging both intra- and intercellular correlations between methylation states. Melissa [14] first defines specific regions of interest in the genome (such as a specific promoter region), then performs generalized linear model regression on CpG sites in that region. The model leverages information from other cells through a shared prior distribution determined by a Bayesian mixture model, effectively clustering cells. DeepCpG [15] proposes a recurrent neural network (RNN) to process differences in local CpG profiles across cells. For every cell, the local CpG profile consists of a vector containing the methylation states and distances of the 25 nearest observed CpG sites up- and downstream from the target site. Along with this RNN, a convolutional neural network (CNN) processes relevant information in the DNA sequence surrounding the target site. The two streams of information are combined near the end of the network. Finally, the output head returns predictions for every cell at a single CpG site. Using similar design principles, LightCpG uses gradient boosting to obtain faster training times at the cost of a lower performance [16]. CaMelia [17], also relying on gradient boosting models, restricts its imputation to CpG sites that are also recorded in at least one other cell. It additionally discards CpG sites whose local methylation profiles are too dissimilar of the profiles in all other cells. Uniquely, CaMelia introduces the notion of using bulk tissue samples to improve performance compared to DeepCpG and trains a separate model for every cell. It remains unclear, however, whether these performance gains can be attributed to the employed methods or to the aforementioned sample selection.

In this work, inspiration is drawn from recent developments in self-supervised learning of natural language. In particular, the language model BERT is trained by randomly replacing words in a sentence by a unique [MASK] token and attempting to predict the masked word given the newly-formed sentence (called masked language modeling) [18]. In essence, this objective trains a model to fill in the gaps in a sentence. The similarity with imputation, where gaps in a matrix need to be filled in, is compelling but unexplored. In language modeling, transformer neural networks are used because of their capability of learning interactions between all input words, akin to the flow of information in a complete digraph [19].

Biological systems can be elegantly represented by graphs [20]. For example, the interactions of genes form distinct pathways in a regulatory network. Consequently, models should ideally reason over graphs or mimic graph structure. Most of the current deep learning practices in bioinformatics do not reflect this reality. For example, fully-connected layers learn a set of fixed weights for all inputs and are hence unable to reason over how correlations between inputs differ when their contents change. Transformers mimic graph structure using a self-attention mechanism to explicitly reason over how every input is influenced by the others [21]. Because of this, transformers scale quadratically in computational- and memory cost with the number of inputs. They have previously been shown to outperform other neural architectures in DNA sequence annotation tasks [22] and protein representation learning [23, 24]. Recently, the use of transformers in biology has gone beyond 1D sequences. For example, MSA Transformer [25] adapts axial attention [26] to MSAs for unsupervised protein structure learning. AlphaFold2 [27] also adapts axial attention to process both MSAs and residue pair matrices. By processing 2D inputs, full self-attention learns 𝒪(*n*^2^*m*^2^) pairwise interactions for a ℝ^*n×m*^ matrix, making vanilla transformers impossible to apply on high-dimensional methylation data.

We introduce CpG Transformer, an adaptation of the transformer neural network architecture to operate on partially-observed methylation matrices by combining axial attention [26] with sliding window self-attention [28], thereby obtaining state-of-the-art imputation performances on a wide range of datasets. The inputs to CpG Transformer consist of the CpG matrix along with their respective positions on the genome and the DNA sequences surrounding them. Cell identity is communicated to the model through learned cell embeddings. The model learns a representation for every CpG site and recombines their information in a graph-like manner. Because of this, the architecture captures general-purpose representations, allowing for quick transfer learning of imputation models on new datasets, a prospect of great interest to practitioners with limited computational resources. In addition, ablation studies and model interpretation demonstrate the contributing factors to single-cell DNA methylation.

## 2 Methods

Here, CpG Transformer is described for the imputation of DNA methylation data. Our architectural contributions are twofold. Firstly, CpG Transformer draws inspiration from collaborative filtering approaches to formulate its inputs to the transformer layers [29]. The transformer layers model the interactions between matrix entries. In this sense, CpG Transformer can be regarded as *contextualized collaborative filtering*. Secondly, we extend axial attention [26] to incorporate sliding window self-attention [28], where full self-attention is applied per individual column and sliding window self-attention is applied over all rows separately.

### Model Inputs

The input to CpG Transformer is a three-dimensional tensor 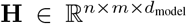, where 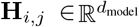 represents the input representation at cell *i* (rows) and methylation site *j* (column) of the methylation matrix. Every representation **H**_*i,j*_ is the result of linear combination of a concatenation of three embeddings: 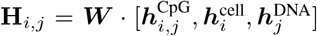 (Figure 1). All three embeddings consist of 32 hidden dimensions, and are combined to *d*_model_ = 64 dimensions by ***W***. The CpG embedding 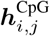 is obtained by embedding the methylation state (unknown ?, unmethylated 0 or methylated 1) of CpG site *j* in cell *i*. Similarly, row-wise cell embeddings 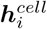 encode a hidden representation for cell indices. Finally, DNA sequence information is included in the model by taking 1001 nucleotide windows centered around the methylation sites and processing them with a CNN to obtain column-wise DNA embeddings 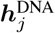. In all experiments, the CNN architecture is adapted from DeepCpG, consisting of 2 convolutional layers, each followed by a max-pooling layer [15]. The exact parameters of the CNN backbone are elaborated in Supplementary Section S1.

**Figure 1:**
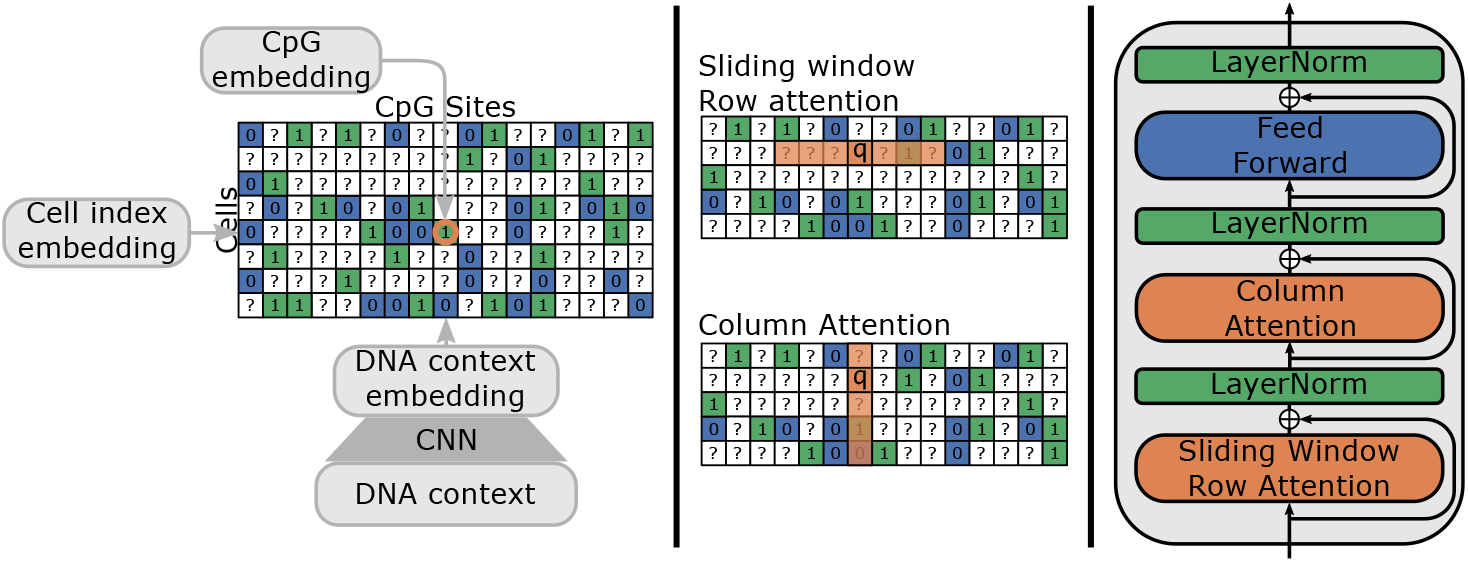
**Left:** Inputs to CpG Transformer. Cell, DNA, and CpG embeddings are applied row-, column-, and element-wise, respectively. **Middle:** Illustration of sliding window row attention and (full) column attention. For sliding window row attention, every query attends to keys in the same row within a fixed window. For column attention, every query attends to the keys from all elements in the same column. **Right:** A single CpG Transformer layer.

### CpG Transformer

Transformer layers employ self-attention to explicitly reason over how every input is influenced by the others [21]. All *n* entries of an input ***X*** are once encoded as a query and once as a key via learned linear layers. For this model setup, the input to the transformer layers is **H**. Taking the inner product of the queries ***Q*** with the keys ***K*** results in an *n* × *n* matrix, whose values can be loosely interpreted as the importance of input *j* for input *i*, at row *I* and column *j*. These values are normalized and multiplied by a value matrix ***V*** (obtained via linear combination of the input ***X*** with learned weights) to produce outputs for every input entry in a matrix ***Z***. This process can be performed multiple times in parallel using separate weight matrices, constituting different attention heads. Corresponding outputs ***Z*** for every head can then be concatenated and linearly combined to an appropriate hidden dimension size. The scaled dot-product self-attention mechanism first described by Vaswani et al. [21] is given by the following equations, where *d*_*k*_ denotes the hidden dimensionality of the queries and keys:

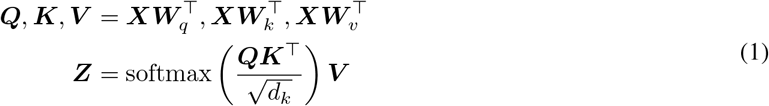

Intuitively, this mechanism simply learns how inputs should be recombined in order to propagate to an output at every position. As such, no structure in the input is assumed and fixed-length inputs are not required, as identical model weights are used for every position. For an input methylation matrix with *n* cells (rows) and *m* methylation sites (columns), an *n* · *m* × *n* · *m* attention matrix would be obtained. Because *m* can easily exceed millions, it is impossible to apply vanilla transformers to methylation data. To reduce the computational complexity of this operation, axial attention [26] can be employed. In axial attention, dependencies between elements of the same row and elements of the same column are modeled separately by two distinct self-attention operations in every layer (Figure 1). In doing so, the memory and computational complexity is reduced from 𝒪(*n*^2^*m*^2^) to 𝒪(*mn*(*n* + *m*)). Further, known autocorrelation between neighboring methylation sites can be leveraged. It is known that CpG sites in close proximity of each other on the genome are often correlated [30]. Hence, we can limit row-wise attention to interactions between neighboring CpG sites in a sliding window attention mechanism [28, 31], further reducing the complexity from 𝒪(*mn*(*n* + *m*)) to 𝒪(*mn*(*n* + *w*)), with a window size *w* (analogous to kernel size in convolutions). In order to communicate relative distances of CpG sites to the model, row-wise sliding window self-attention operation is supplied with relative sinusoidal positional encodings [32]. Pseudocode describing both row- and column-wise self-attention operations in more detail can be found in Supplementary Section S2.

CpG Transformer employs a stack of four identical layers (Figure 1). The layer structure is similar to the one defined by Vaswani et al. [21] and Rao et al. [25]. Each layer has three sub-layers. The first and second sub-layers consist of the previously-described column-wise self-attention and row-wise sliding window self-attention with 8 heads of 8 hidden dimensions each. A window size of *w* = 41 is used for the sliding window self-attention (analogous to a convolutional kernel size of 41). The window size is selected considering a trade-off between computational complexity and inclusion of biological information. A larger window size means that CpG Transformer recombines information from more neighboring sites at the cost of computational- and memory complexity. The input and output dimensionality of the attention layer is *d*_model_ = 64. The last sub-layer employs a position-wise fully-connected feed-forward network consisting of two linear combinations with a ReLU activation in between: max(0, ***XW***_1_ + ***b***_1_)***W***_2_ + ***b***_2_. The dimensionality of input and output is *d*_model_ = 64 and the inner-layer has 256 hidden dimensions. A residual connection [33] followed by layer normalization [34] is employed around all sub-layers. The outputs of the last transformer layer are reduced to one hidden dimension by an output head and subjected to a sigmoid operation to obtain final predictions ***Ŷ*** ∈ ℝ^*n×m*^ for all inputs.

### Training Objective

We adapt the masked language modeling (MLM) objective for DNA methylation imputation [18]. MLM is a type of denoising autoencoding in which the loss function acts only on the subset of inputs that are perturbed. For CpG Transformer, the inputs are corrupted by randomly masking observed sites to the ? token. In addition, 20% of the tokens that would be masked are instead randomized to a random state (0 or 1), sampled proportionally to the distribution of methylation states in the input. In doing so, CpG Transformer learns not only to impute but also to denoise. Finally, the cross-entropy loss optimizes the model to return the original methylation states given the corrupted input. Devlin et al. [18] additionally proposes to leave a percentage of the masked tokens to be unchanged instead. Considering our limited vocabulary size (unknown ?, unmethylated 0 or methylated 1), a sufficient percentage of randomized tokens is actually unchanged, eliminating the need for this operation. An overview of the training procedure is given in Figure 2.

**Figure 2:**
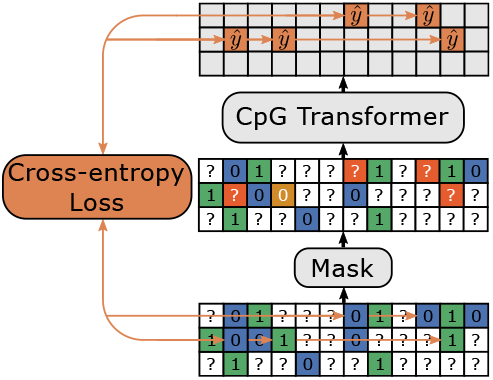
Masked language modeling. Positive and negative sites are indicated in green and blue, respectively. Sites to train on (orange) are either masked (80%) or randomized (20%). The model is optimized to infer the original methylation state given the corrupted input using the cross-entropy loss.

### Datasets

Five publicly available datasets originating from both scBS-seq [4] and scRRBS-seq [5] experiments are obtained from the Gene Expression Omnibus.

The first dataset (GSE56879) consists of 20 mouse embryonic stem cells cultured in Serum. The second dataset is obtained from the same study and is made up of 12 cells of the same type cultured in 2i medium [4]. Both datasets were profiled using scBS-seq. A third dataset (GSE65364) comprises 25 human hepatocellular carcinoma cells profiled using scRRBS-seq [35]. scRRBS-seq profiles of 30 human monoclonal B-cell lymphocytes form a fourth dataset (GSE125499; sc05) [36]. The final dataset (GSE87197) consists of 122 hematopoietic stem cells and progenitor cells profiled using scBS-seq [7]. This dataset includes 18 hematopoietic stem cells, 18 multipotent progenitors, 19 common myeloid progenitors, 24 multi-lymphoid progenitors, 22 granulocyte macrophage progenitors and 21 common lymphoid progenitors. In the remainder of this paper, these datasets are referred to as Ser, 2i, HCC, MBL, and Hemato, respectively. Corresponding reference genomes are as follows: Ser and 2i use genome build NCBIM37. GRCh38 is used by Hemato, and GRCh37 serves as reference genome for HCC and MBL. A brief summary of dataset sizes is available in Supplementary Table S1.

For all datasets, binary methylation states are obtained by assigning a positive (methylated) label when 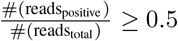. We use holdout validation to test the performance of the models. For all datasets and experiments, chromosome 5 and 10 constitute the validation and test set, respectively. All other chromosomes are used in training. More instructions on how to obtain and preprocess the datasets, as well as their corresponding reference genomes, are available on the GitHub page of CpG Transformer.

### Models and Training

CpG Transformer is compared to two competing methods: DeepCpG [15] and CaMelia [17]. A comparison with Melissa is not considered since the method is described as complementary to whole-genome imputation methods [14]. For all datasets, the default hyperparameters of CaMelia and DeepCpG are used. To ensure a fair comparison, all models are trained using the same data preprocessing and splits. Due to this, performances are expected to deviate slightly from those reported in their respective manuscripts. Considering reproducibility concerns, a full list of differences in our implementations of DeepCpG and CaMelia is given in Supplementary Section S1.

A separate CpG Transformer with identical hyperparameters is trained for every dataset on 2 V100 GPUs using Adam as optimizer [37]. A learning rate of 5 · 10^−4^ with linear warmup over the first 1000 steps is used. The learning rate is multiplicatively decayed by a factor 0.9 after every epoch. Models are trained for a maximum of 100 epochs, with early stopping after no validation loss decrease has been observed for 10 epochs. The model arising from the epoch with the best validation loss is kept as final model. A dropout rate of 0.20 on elements of the attention matrix is employed during training. Batches are constructed by slicing the *n* × *m* methylation matrices vertically into *n* × *b* bins with *b* = 1024 CpG sites each. One such a bin makes up a batch. For every batch, the number of sites that are masked or randomized equals 25% the number of columns in the bin for all datasets. This masking percentage is chosen considering that a large proportion of the input already consists of masked ? tokens. For the Hemato dataset, we additionally randomly subsample 32 rows (cells) every training batch to reduce complexity and increase training speed. Finally, because random masking negatively biases evaluation, test performance is measured by masking every methylation site in the dataset separately in smaller batches (Note that this is only necessary to fairly compare imputation performance on all available labels. In practice, inference would be performed without masking).

## 3 Results

### 3.1 Imputation performance

To benchmark CpG Transformer, we evaluate against one competing deep learning method, DeepCpG [15], and one traditional machine learning method, CaMelia [17]. The resulting imputation performances in terms of area under the receiver operating characteristic curve (ROC AUC) and area under the precision-recall curve (PR AUC) for all datasets are shown in Table 1. CpG Transformer consistently outperforms existing models on all datasets. In addition, CpG Transformer uses a similar amount of computational budget as DeepCpG and CaMelia (Supplementary Table S2). Cell-specific performance evaluation (Supplementary Figure S1) shows that CpG Transformer is, out of all cells, only outperformed by competing methods for a single Hemato cell. Furthermore, CpG Transformer consistently outperforms DeepCpG and CaMelia in a variety of genomic contexts (Supplementary Figure S2). The performance gain is most pronounced in contexts typically associated with higher cell-to-cell variability, such as CpG islands, regulatory elements and histone modification marks [38], demonstrating CpG Transformer’s ability to encode relevant cell heterogeneity.

**Table 1:** Performance comparison of CpG Transformer with other methods. Sparsity is defined as the percentage of entries in the methylation matrix that are unobserved. Best performers are indicated in bold. The reported metrics are computed for all cells together.

| Dataset | # Cells | Sparsity (%) | ROC AUC |  |  | PR AUC |  |  |
| --- | --- | --- | --- | --- | --- | --- | --- | --- |
|  |  |  | DeepCpG | CaMelia | CpG Transformer | DeepCpG | CaMelia | CpG Transformer |
| Ser | 20 | 77.8 | 90.21 | 90.22 | **91.55** | 92.77 | 92.86 | **93.87** |
| 2i | 12 | 77.9 | 84.80 | 83.02 | **85.77** | 71.69 | 68.87 | **73.56** |
| HCC | 25 | 88.5 | 96.89 | 97.42 | **97.96** | 92.58 | 94.10 | **95.19** |
| MBL | 30 | 90.8 | 88.22 | 89.17 | **92.49** | 87.61 | 87.6 | **91.80** |
| Hemato | 122 | 98.4 | 88.85 | 89.16 | **90.65** | 95.60 | 95.84 | **96.43** |

A small ablation study (Table 2) on the Ser dataset shows the importance of the different inputs to the model. The original model is compared to four models, each trained and evaluated in a scenario where one specific input is left out: 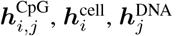, or the positional encodings. Without the CpG embedding, the model can only rely on cell identity and the DNA contexts of their own and neighboring CpG sites. The model without this embedding displays the lowest performance, illustrating the key importance of dependencies between methylation states for their prediction. Without cell embeddings, cell identity is lost and the prediction for every site is the same for all cells. As the second-most important input for the Ser dataset, this embedding highlights CpG Transformer’s capability to exploit cell heterogeneity. Without positional encodings, the model has no way of knowing how far away two CpG sites are from each other. Since column-wise correlation between CpG sites decreases with distance [30], their role is to inform the effect of genomic distance on the degree of correlation in a flexible way. In practice, a minimal but noticeable effect of this encoding on the Ser dataset is observed. Consistent with the findings of DeepCpG [15], the DNA embeddings, informing the model of DNA context surrounding CpGs, is indicated as the least important input for imputation of the Ser dataset. Further ablation studies of CpG Transformer hyperparameters (Supplementary Table S3 and Supplementary Table S4) show that scaling the architecture of CpG Transformer up or down does not significantly increase performance.

**Table 2:**
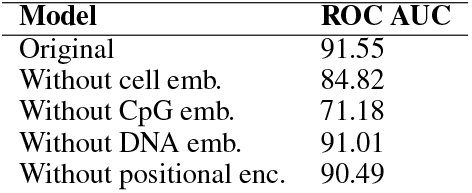
Ablation study on Ser dataset. The original model is compared to four models for which one type of input is removed.

| Model | ROC AUC |
| --- | --- |
| Original | 91.55 |
| Without cell emb. | 84.82 |
| Without CpG emb. | 71.18 |
| Without DNA emb. | 91.01 |
| Without positional enc. | 90.49 |

The performance of all models is heavily dataset-dependent, indicating their varying quality. Since single-cell sequencing experiments suffer from low sequencing depth, we hypothesize that performance is negatively influenced by limited coverage both at the CpG site in question (noisy labels) and in its neighborhood (in terms of number of unobserved entries, termed local sparsity). To test this, the performance of the Ser dataset in function of these factors is plotted (Figure 3). Similar plots for the other datasets are shown in Supplementary Figure S3. It is observed that CpG sites covered by a smaller number of reads have a less-confident label, resulting in negatively-biased performance at evaluation. In addition, CpG sites with a higher local sparsity are harder to predict, presumably due to providing a noisier estimate of local methylation profiles. By making a heatmap of performance in function of both these factors, it is observed that local sparsity is most causal of lower predictive performance. In the context of these experiments, we note that a perfect imputation performance is realistically unattainable given the inherent noise in single-cell methylation datasets.

**Figure 3:**
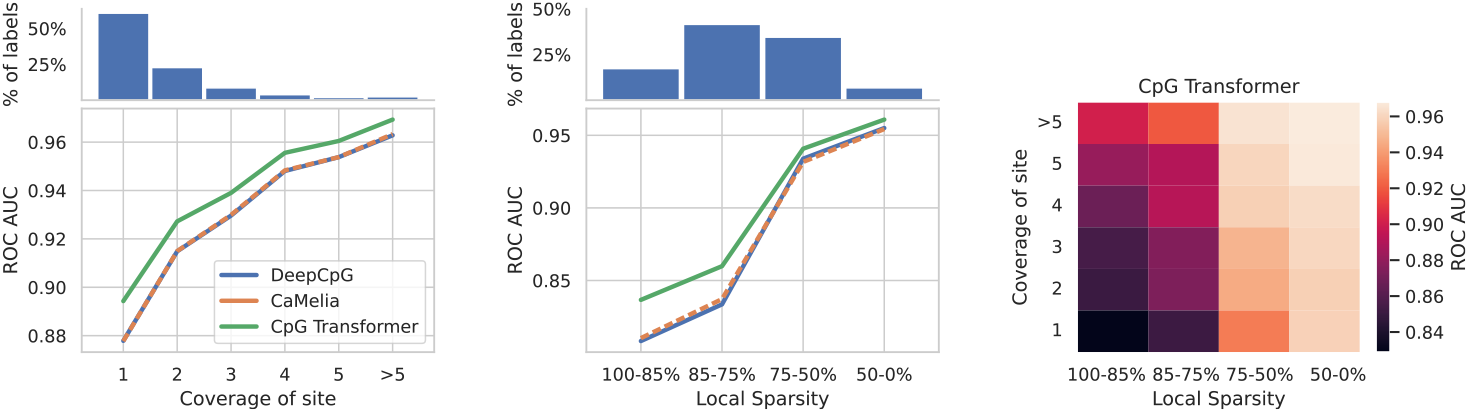
Dependency of performance on sequencing depth for the Ser dataset. **Left:** ROC AUC in function of coverage (# reads) of the label. On top of the plot the percentage of labels belonging to each bin is shown. **Middle:** ROC AUC in function of local sparsity (defined as the percentage of unobserved entries in the local window used for prediction). **Right:** ROC AUC in function of both factors for CpG Transformer. The biggest gradient in performance is observed for the local sparsity direction.

### 3.2 Model interpretation

Because CpG Transformer recombines information from CpG sites in a general way, it lends itself well to model interpretation methods. Here, we aim to attribute model predictions to its input features, a problem best approach with gradient-based saliency methods [39]. Integrated Gradients [40] computes the gradients of the prediction with respect to the input features to measure how every input contributes to prediction. Contributions are obtained by decomposing the difference in prediction of the input sample with an all-zero baseline. For CpG Transformer, contribution scores are obtained for all inputs to the first transformer layer. Since four transformer layers with a window size of 41 are employed, the total receptive field for any prediction constitutes the 161 surrounding CpG sites for all *n* cells (*n* × 161). Because Integrated Gradients returns contribution scores for all hidden dimensions, they are summed to obtain a single score for every input matrix entry. An example contribution for the Ser dataset is shown in Figure 4A.

**Figure 4:**
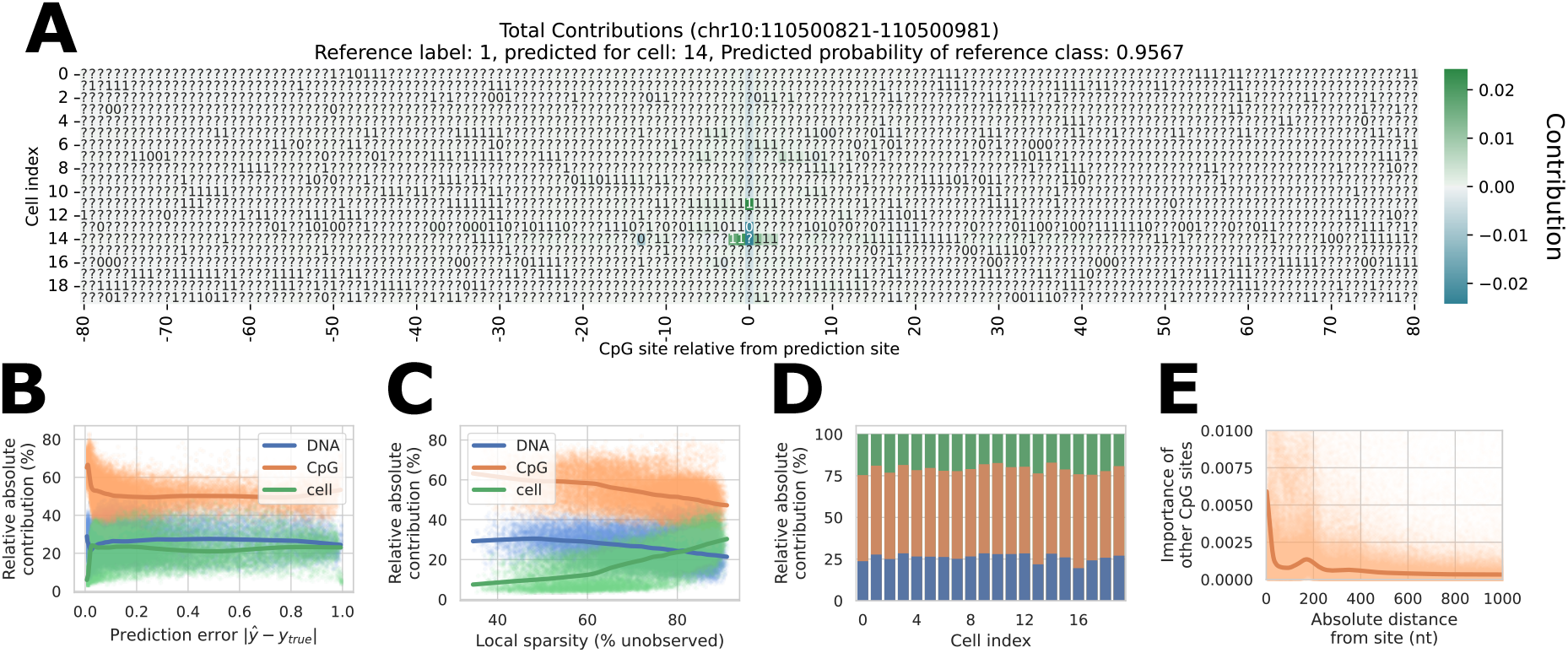
Integrated Gradients interpretation for the Ser dataset. All trend lines are obtained with LOWESS. [**41**]. **A:** Example Integrated Gradients contribution plot. The contributions for the prediction of a randomly chosen CpG site in cell 14 are shown The true label of the site was originally 1, but is masked in the input here. Matrix entries with a positive and negative contribution are colored in green and blue, respectively. **B:** Relative absolute contribution of the three input embeddings in function of prediction error. **C:** Relative absolute contribution in function of local sparsity. **D:** Relative absolute contribution stratified per cell. **E:** Contribution of observed CpG sites in function of their absolute distance to the prediction site.

The contributions of the individual matrix entries can be decomposed into those of their constituent embeddings 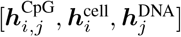 by backpropagating Integrated Gradients one layer further. In doing so, contribution matrices similar to the one shown in Figure 4A are obtained for all three embeddings (Supplementary Figure S4). Performing this for 1% of the samples in the test set, it is possible to investigate how embeddings contribute to prediction in different settings. This way, the total contribution of the embeddings in function of the prediction error, local sparsity and cells are obtained for the Ser dataset 4B-D. The same plots for the other datasets are shown in Supplementary Figure S5. We find that CpG embeddings relatively contribute more to predictions when the model is confident (with a small prediction error). In cases where the local sparsity is low (i.e. a low number of unobserved sites), CpG Transformer can rely more on local methylation profiles to make a prediction, increasing the relative importance of CpG embeddings. Between different cells, relative contribution differences are negligible. Figure 4E shows the contributions of neighboring observed CpG sites in function of their distance from the prediction site. A decreasing trend is observed with distance, with one bell-shaped bump appearing at ±160 nucleotides from the prediction site. This relation has been reported on the same cell types in literature by Song et al. [42], who suggested a relation between nucleosome modifications and DNA methylation.

### 3.3 Transfer learning

Because of the generality of CpG Transformer’s self-attention mechanism, it is expected that it learns general-purpose representations of DNA methylation dynamics. In this respect, CpG Transformer is envisioned to transfer well to other datasets. In this paper, transfer learning is examined in the context of improved convergence speed when fine-tuning a trained imputation model to impute a new dataset, a prospect of great interest to practitioners with a limited time and computational budget.

As an experiment, the dynamics of models learning to impute the HCC dataset are investigated (Figure 5). Two CpG Transformer models are trained in the same way as in previous experiments: once with weights initialized randomly as before and once with weights initialized from the model trained on the MBL dataset. All model weights apart from the cell embeddings are transferred. Both training modes are run in triplicate.

**Figure 5:**
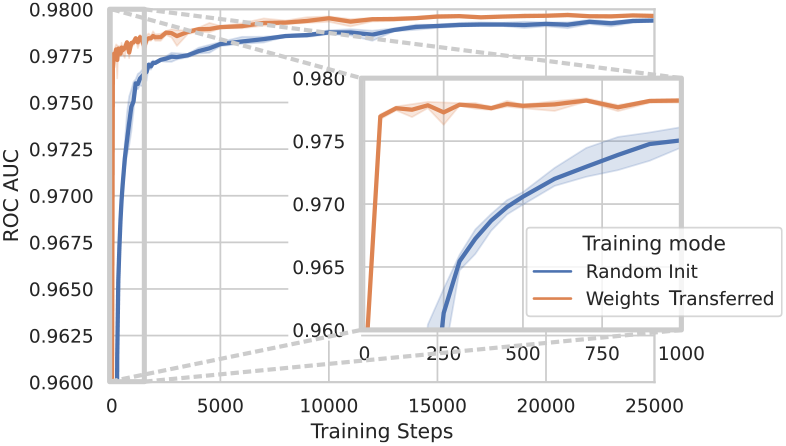
Transfer learning dynamics on the HCC dataset. Two types of models are trained to impute the HCC dataset: once from a random initialization of weights and once with weights initialized from the model trained on the MBL dataset. Error bands indicate the standard deviation of performances over three runs.

The highest achieved performance of both models is similar (97.96 and 97.98 ROC AUC for random and transferred, respectively), but the models with transferred weights converge substantially faster, reaching a ROC AUC within 0.5% of the best performance after only 50 training steps, whereas the randomly initialized model needs 1000 training steps to reach the same performance, indicating an approximate convergence speed up of x20. Furthermore, without any fine-tuning steps, transferred models still achieve a ROC AUC of 95.62, indicating the ability of the transformer weights to figure out cell identity from random cell embeddings. Together, these results show CpG Transformer is accessible to researchers wanting to train an imputation model on their own dataset with a limited computational budget.

## 4 Discussion

CpG Transformer adapts the transformer architecture to operate directly on methylation matrices by combining axial attention [26] with sliding window attention [28], providing a general-purpose way of learning interactions between neighboring CpG sites both within- and between cells. This approach gives rise to many advantages over competing methods. Most simple of all, state-of-the-art imputation performances are obtained over DeepCpG [15] and CaMelia [17]. Secondly, our method lends itself well to interpretation and transfer learning. Finally, because CpG Transformer’s model architecture uses learned cell embeddings to encode cell identity in a flexible way, we envision CpG Transformer to scale better to future larger datasets containing diverse cell types.

CpG Transformer allows the prediction of methylation states of thousands of CpG sites in parallel. It does, however, scale quadratically with the number of cells in the dataset. Given the size of the datasets used in this study, this did not pose a problem. For datasets consisting of thousands of cells, however, the application of CpG Transformer as outlined here becomes impossible. In this case, practitioners would need to split their dataset in multiple smaller subsets in which cells are as similar as possible. Alternatively, further extensions of the proposed axial attention could be made in order to allow inputs with a large number of cells. Self-attention sparsity for the column-wise attention operation could, for example, be enforced through clustered attention [43]. In doing so, interactions would only be modeled between clustered, closely-related cells, instead of between all cells. We consider such extensions to be future work.

The proposed axial attention attends to neighboring sites within a fixed window, irregardless of whether these neighbors have an observed label or not. A possible disadvantage of this strategy may be that, in cases with extreme sparsity, the model may not be able to properly estimate local methylation profiles. In this case, one approach would be not to model interactions within a fixed local window, but instead to attend to the *n* nearest neighboring observed entries in every cell. This mechanism would attend to a fixed number of observed CpG sites independent of local sparsity. Since such a mechanism would attend to sites far away on the genome in high sparsity settings, its added value is not straightforwardly estimated. Another approach would be to attend only to CpG sites within a fixed genomic width (e.g. 1kbp). Unlike the previous proposed mechanism, this method would be at an advantage or disadvantage in regions with high or low CpG density, respectively. We consider comparisons with these approaches to be future work.

Model analysis and interpretation show that local sparsity is an obstacle for the performance of imputation models. Figure 3 surprisingly shows that lowly-covered sites (whose labels are expected to be more noisy) can be more accurately predicted in a densely-covered neighborhood. Some nuances should be made regarding genomic regions that are densely covered but only by a small number of reads for every site. CpG Transformer’s masking and randomizing objective (falsely) assumes no structure in noise and missingness. In reality, for example, one read covering two neighboring unmethylated sites could falsely report methylated signal for both sites if bisulfite treatment failed to convert the corresponding sequence. Hence, lowly-covered sites in densely-covered neighborhoods may be collectively noisy in the same, non-random way. Most contemporary imputation methods, including CpG Transformer, have no way of coping with systematic noise and missingness. In these cases, models will most likely propagate and amplify the noise, potentially compromising biologically-relevant results.

Notwithstanding the above-mentioned considerations, given careful evaluation, CpG Transformer can greatly enhance single-cell methylation studies. A cautious practitioner may, for example, wish to only retain imputations in regions where local sparsity is low and coverage of labels is high. To aid researchers in understanding their imputation results, interpretation methods are introduced. In addition, transfer learning experiments show that CpG Transformer can be used to obtain state-of-the-art imputation performances on a limited time and computational budget.

## Data Availability

The data underlying this article is freely available from the Gene Expression Omnibus following the identifiers listed in Section 2. Source code for CpG Transformer is freely available at https://github.com/gdewael/cpg-transformer.

