## Supplementary Information for "CpG Transformer for imputation of single-cell methylomes"

#### 1 Differences between original and our implementation of DeepCpG and CaMelia

**DeepCpG** We:

- Trained the model in an end-to-end fashion instead of training the CNN and RNN modules separately, as no apparent difference in performance could be found.
- Used the same model hyperparameters for all datasets:
  DNA module = conv[128@11]_mp[4]_conv[256@3]_mp[2]_fc[128]_do[0.0] **(This same convolutional architecture was used in CpG Transformer)**
  CpG module = fc[256]_bgru[256]_do[0.0]
  Joint module = fc[512]_do[0.0]_fc[512]_do[0.0]
- Encoded labels as positive (methylated) label when $\frac{\#(reads_{positive})}{\#(reads_{total})} \geq 0.5$ and otherwise as negative (instead of using $\frac{\#(reads_{positive})}{\#(reads_{total})} > 0.5$), as slightly better performance could be obtained using this encoding scheme.
- Encoded nucleotides with an embedding layer instead of one-hot encoding.
- Used different chromosomes in holdout validation and testing.
- Used all cells for the Ser dataset.
- Improved the efficiency of preprocessing: batches are constructed on the fly without having to precompute the neighbors and write them to disk.
- Used linear learning rate warm-up over the first 1000 steps, through which more-stable learning trajectories are obtained.

**CaMelia** We:

- Encoded genomic positions of CpG sites as 64-bit integers instead of 32-bit floats.
- Excluded bulk cells from the analysis.
- Did not discard samples with an even number of methylated and unmethylated reads. Instead, labels are encoded as positive (methylated) label when $\frac{\#(reads_{positive})}{\#(reads_{total})} \geq 0.5$.
- Did not rule out samples which originate from CpG sites that do not have a label in any other cell.
- Did not discard samples for which no other cell has a local similarity of larger than 0.8. CaMelia excludes these samples because the locally paired similarity feature is undefined in these cases. In order to obtain a prediction for every methylation state in every cell (as is possible with CpG Transformer and DeepCpG), the locally paired similarity feature for these problematic samples is instead assigned with `NaN`. The default processing mode for `NaN` values in CatBoost is processing them as the minimum value for that feature.
- Used different chromosomes in holdout validation and testing, instead of using cross-validation.
- Improved the efficiency of preprocessing: locally paired similarity features are computed in a vectorized way wherever possible. Feature vectors are computed and encoded in a memory-efficient way, so that they do not have to be written to disk.

#### 2 Pseudocode for self-attention operations

**Algorithm 1:** Column-wise full self-attention

```
# b = batch size, n = number of rows/cells, m = number of columns/CpGsites, d_model = hidden dimensionality
Data: X ∈ ℝ^{b×n×m×d_model}
Result: Z ∈ ℝ^{b×n×m×d_model}
Q ← MatMul(X, W_q) ∈ ℝ^{b×n×m×(n_head d_head)} # n_head = number of heads, d_head = dims per head
K ← MatMul(X, W_k) ∈ ℝ^{b×n×m×(n_head d_head)}
V ← MatMul(X, W_v) ∈ ℝ^{b×n×m×(n_head d_head)}
# To perform attention separately for each column:  batch and column dimension are collapsed.
# In addition, attention heads are splitted in a separate dimension.
Q ← Reshape(Q) ∈ ℝ^{(bm)×n×n_head×d_head}
K ← Reshape(K) ∈ ℝ^{(bm)×n×n_head×d_head}
V ← Reshape(V) ∈ ℝ^{(bm)×n×n_head×d_head}
# Scaled dot-product self-attention.
A ← MatMul(Q, K^⊤)/√d_head ∈ ℝ^{(bm)×n×n×n_head}
A ← Dropout(Softmax(A))
Z ← MatMul(A, V) ∈ ℝ^{(bm)×n×n_head×d_head}
# Disentangling batch and column dimension again.
Z ← Reshape(Z) ∈ ℝ^{b×n×m×(n_head d_head)}
Z ← Dense(Z) ∈ ℝ^{b×n×m×d_model}
```

**Algorithm 2:** Row-wise sliding-window self-attention, with positional encodings as in Transformer-XL.

```
# b = batch size, n = number of rows/cells, m = number of columns/CpGsites, d_model = hidden dimensionality
Data: X ∈ ℝ^{b×n×m×d_model}, Positional encoding: P ∈ ℝ^{(bn)×m×n_head×d_head}, trainable bias vectors: u ∈ ℝ^{d_head} and
      v ∈ ℝ^{d_head}
Result: Z ∈ ℝ^{b×n×m×d_model}
Q ← MatMul(X, W_q) ∈ ℝ^{b×n×m×(n_head d_head)} # n_head = number of heads, d_head = dims per head
K ← MatMul(X, W_k) ∈ ℝ^{b×n×m×(n_head d_head)}
V ← MatMul(X, W_v) ∈ ℝ^{b×n×m×(n_head d_head)}
# To perform attention separately for each row:  batch and row dimension are collapsed.
# In addition, attention heads are splitted in a separate dimension.
Q ← Reshape(Q) ∈ ℝ^{(bn)×m×n_head×d_head}
K ← Reshape(K) ∈ ℝ^{(bn)×m×n_head×d_head}
V ← Reshape(V) ∈ ℝ^{(bn)×m×n_head×d_head}
# To perform sliding window self-attention:  m sliding windows of size w are taken from the column dimension.
K ← Unfold(K) ∈ ℝ^{(bn)×m×w×n_head×d_head}
V ← Unfold(V) ∈ ℝ^{(bn)×m×w×n_head×d_head}
# Scaled dot-product self-attention as in Transformer-XL.
A ← [MatMul(Q + u, K^⊤) + MatMul(Q + v, P^⊤)]/√d_head ∈ ℝ^{(bn)×m×w×n_head}
A ← Dropout(Softmax(A))
Z ← MatMul(A, V) ∈ ℝ^{(bn)×m×n_head×d_head}
# Disentangling batch and row dimension again.
Z ← Reshape(Z) ∈ ℝ^{b×n×m×(n_head d_head)}
Z ← Dense(Z) ∈ ℝ^{b×n×m×d_model}
```

### 3 Tables supporting the results section

Table 1: **Summary of dataset statistics.**

| Dataset | # cells (rows) | # CpG sites (columns) | # observed sites |
|---|---|---|---|
| Ser | 20 | 17 315 500 | 76 920 535 |
| 2i | 12 | 15 023 258 | 39 817 398 |
| Hemato | 122 | 18 050 756 | 34 855 325 |
| HCC | 25 | 2 044 635 | 5 891 193 |
| MBL | 30 | 4 779 569 | 13 155 172 |

Table 2: **Comparison of computational budget needed to train CpG Transformer and other models on the HCC dataset.** Training times are measured using a single V100 GPU and 8 cores of an Intel Xeon Gold 6242 (Cascade Lake @ 2.8 GHz). All times are obtained using our optimized reimplementations of all models. Obtained times are represented as the mean ± standard deviation over three runs. DeepCpG trains and performs inference in about half the time as CpG Transformer. CaMelia also trains slightly faster than CpG Transformer, but because it constructs a separate model for every cell, CaMelia takes significantly more time for doing inference and preprocessing. It has to be noted that the HCC dataset, while numbering 25 cells, has the lowest number of CpG sites (columns) of all tested datasets.

| Model | GPU memory during training [a] | # Total weights | # Transformer or RNN weights | Preprocessing time (min) | Training time (min)[b] | Inference time (min) |
|---|---|---|---|---|---|---|
| CpG Transformer | 8299MB [c] | 4.4M | 284K | 0.35 ± 0.01 | 36.12 ± 3.91 | 1.93 ± 0.04 |
| DeepCpG | 1431M | 5.5M | 815K | 1.04 ± 0.02 | 16.21 ± 2.02 | 1.15 ± 0.02 |
| CaMelia | NA | NA | NA | 26.81 ± 0.15 | 19.05 ± 0.38 | 10.93 ± 0.17 |

[a] GPU memory is influenced by both the dataset and hyperparameters. For example, all models should scale linearly with the window size, whereas CpG Transformer scales quadratically with the number of cells (as opposed to linearly for DeepCpG). Note that the same statements on scaling with cells and window sizes can be made for training and inference time.

[b] For CpG Transformer and DeepCpG, training time is defined as the time it takes to reach a ROC AUC within 0.1% of the best performance. For CaMelia, training time is determined by CatBoost default convergence parameters.

[c] Using a default bin size of 1024 CpG sites, as opposed to the batch size of 128 used in DeepCpG. When CpG Transformer is trained with a bin size of 128 CpG sites, 1909MB GPU memory is used.

Table 3: **Ablation study of self-attention mechanisms on Ser dataset.** The final proposed CpG Transformer consists of an axial self-attention mechanism where sliding window row attention is performed first followed by column-wise full self-attention. Full 2D sliding window self-attention refers to a self-attention mechanism where both row-wise sliding window and column-wise self-attention are combined into one operation, as opposed to following each other as in axial attention. Best performers are indicated in bold.

| Model | Sliding window size | ROC AUC | PR AUC |
|---|---|---|---|
| No Self-Attention | / | 86.39 | 89.89 |
| Only Sliding Window Row (intracellular) | 11 | 90.11 | 92.61 |
| Only Sliding Window Row (intracellular) | 21 | 90.19 | 92.65 |
| Only Sliding Window Row (intracellular) | 41 | 90.36 | 92.85 |
| Only Column (intercellular) | / | 89.06 | 92.13 |
| Axial | 11 | 91.44 | 93.78 |
| Axial | 21 | 91.51 | 93.84 |
| Axial | 41 | **91.55** | **93.87** |
| 2D sliding window self-attention | 11 | 91.46 | 93.79 |
| 2D sliding window self-attention | 21 | 91.49 | 93.81 |
| 2D sliding window self-attention | 41 | 91.52 | 93.84 |

Table 4: **Ablation study of hyperparameters on the Ser dataset.** The model is compared to smaller, bigger, deeper and more shallow versions of itself, as well as a version where the order of row-wise and column-wise self-attention is switched. Reported test performances are the mean ± standard deviation over three independent seeds. It can be seen that no considerable performance gains can be made from further scaling up CpG Transformer's weights. The final model (in bold) uses a model size that is as minimal as possible, without negatively inhibiting performance to a considerable extent.

| Model | # transformer weights | # layers | $d_{model}$ | ROC AUC | PR AUC |
|---|---|---|---|---|---|
| CpG Transformer (column-row attention) | 284 160 | 4 | 64 | 91.37 ± 0.03 | 93.56 ± 0.01 |
| CpG Transformer (Small) | 72 448 | 4 | 32 | 91.44 ± 0.07 | 93.83 ± 0.04 |
| CpG Transformer (Big) | 1 125 376 | 4 | 128 | 91.55 ± 0.03 | 93.88 ± 0.01 |
| CpG Transformer (Shallow) | 142 080 | 2 | 64 | 91.49 ± 0.06 | 93.85 ± 0.02 |
| CpG Transformer (Deep) | 426 240 | 6 | 64 | 91.56 ± 0.05 | 93.88 ± 0.01 |
| **CpG Transformer** | **284 160** | **4** | **64** | **91.54 ± 0.04** | **93.87 ± 0.01** |

### 4 Figures supporting the results section

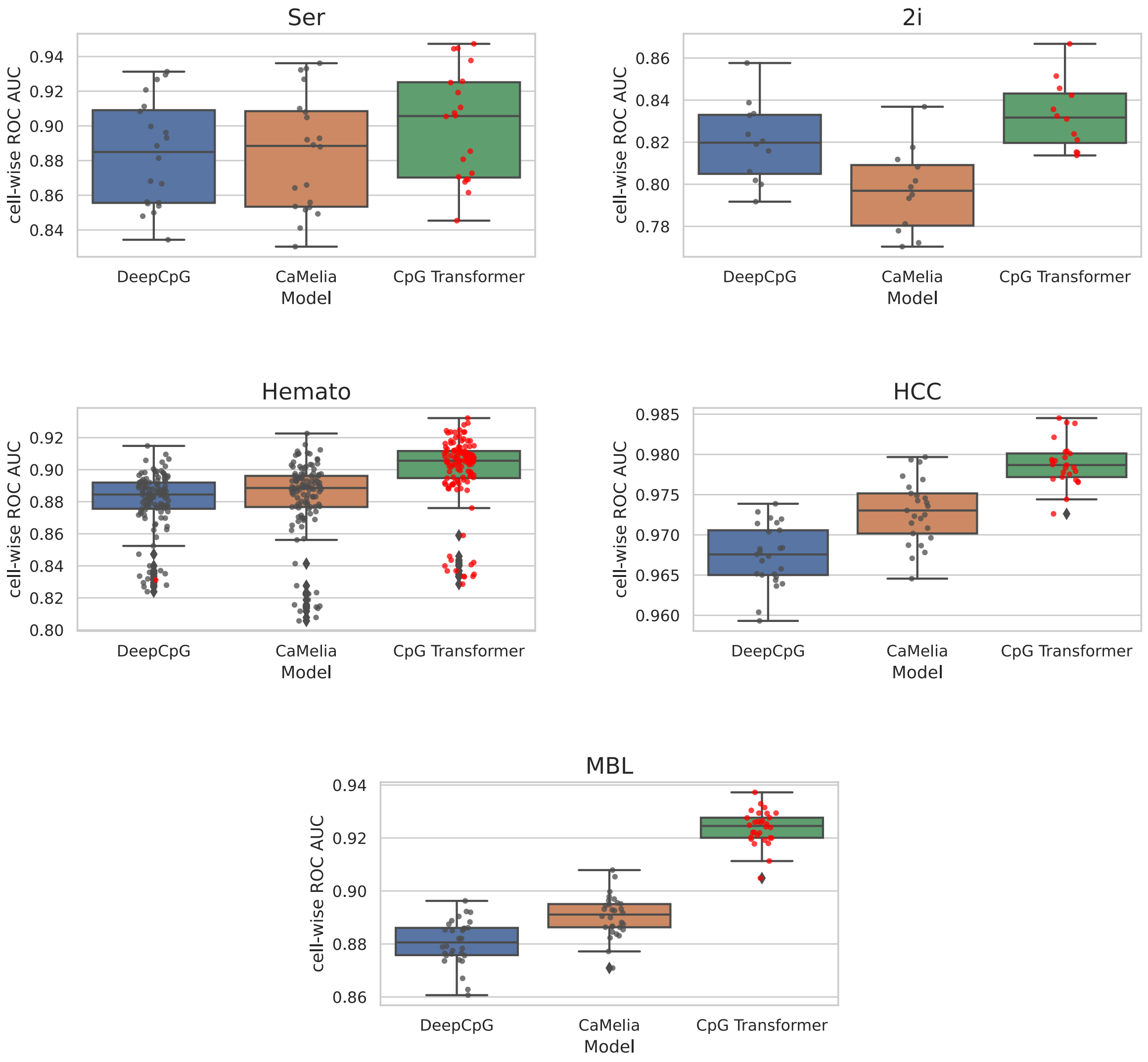


Figure 1: **Boxplots of ROC AUCs per cell.** Every cell is additionally shown by a dot. The best performing model for every cell is indicated by red dots.

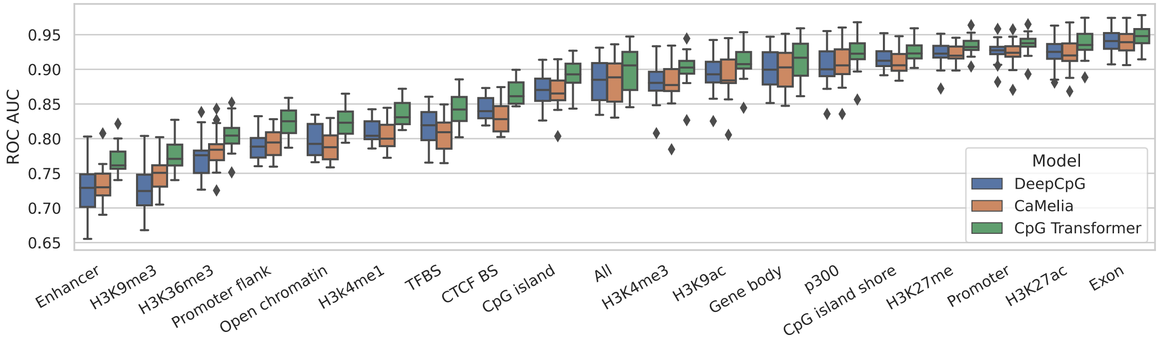


Figure 2: **Boxplots of ROC AUCs per cell on Ser dataset, stratified by genomic contexts.** CpG Transformer outperforms previous models across all tested genomic contexts. Genomic contexts are obtained from Ensembl release 104.

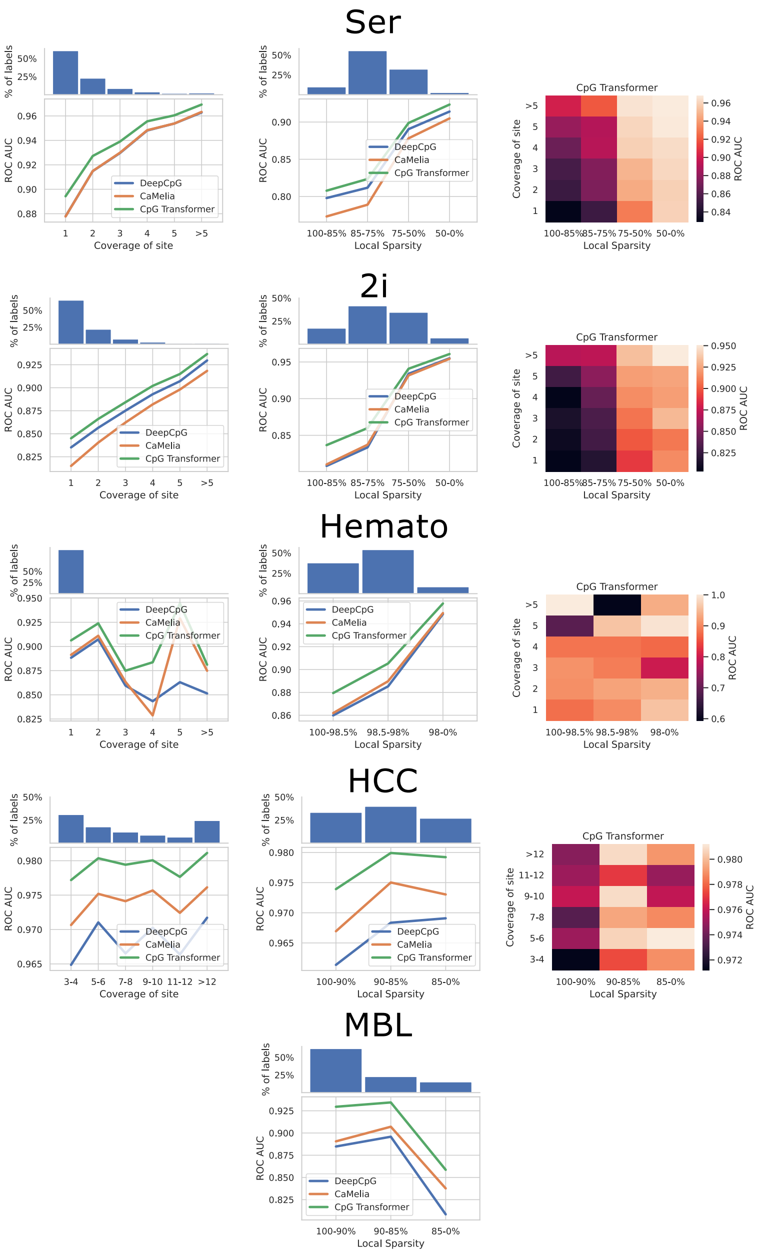


Figure 3: **Dependency of performance on sequencing depth. Left:** ROC AUC in function of coverage (# reads) of the label. On top of the plot the percentage of labels belonging to each bin is shown. **Middle:** ROC AUC in function of local sparsity **Right:** ROC AUC in function of both factors for CpG Transformer. Rows indicate the different datasets. From top to bottom: Ser, 2i, Hemato, HCC and MBL. The biggest gradient in performance is observed for the local sparsity direction. No coverage dependency could be obtained for the MBL dataset as no read data is available for this dataset. Generally, local sparsity is more indicative of low performance than coverage of the label.

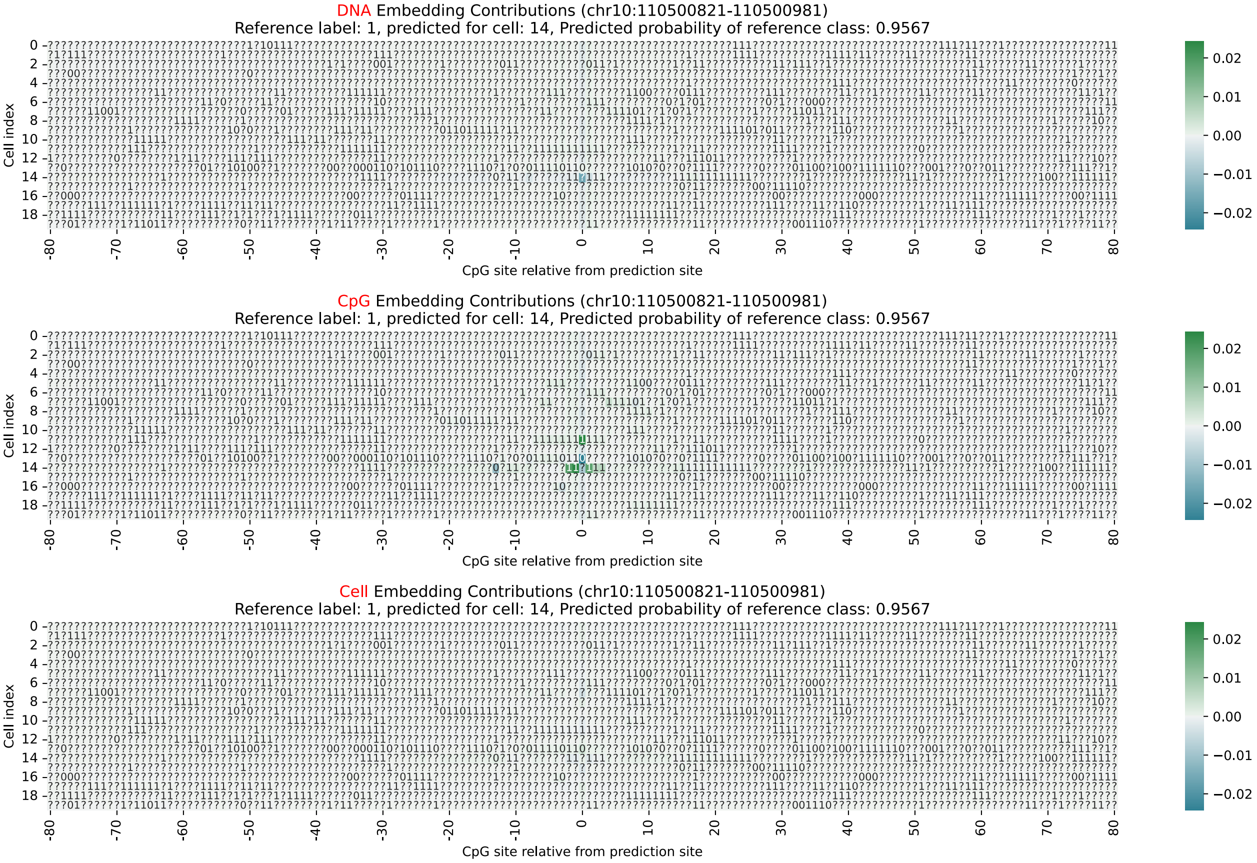


Figure 4: **Example Integrated Gradients Contributions plot.** The contributions for the prediction of a single CpG site in cell 14 are shown. Matrix entries with a positive and negative contribution are colored in green and blue, respectively. The contributions of DNA, CpG and cell embeddings are shown on the top, middle and bottom, respectively.

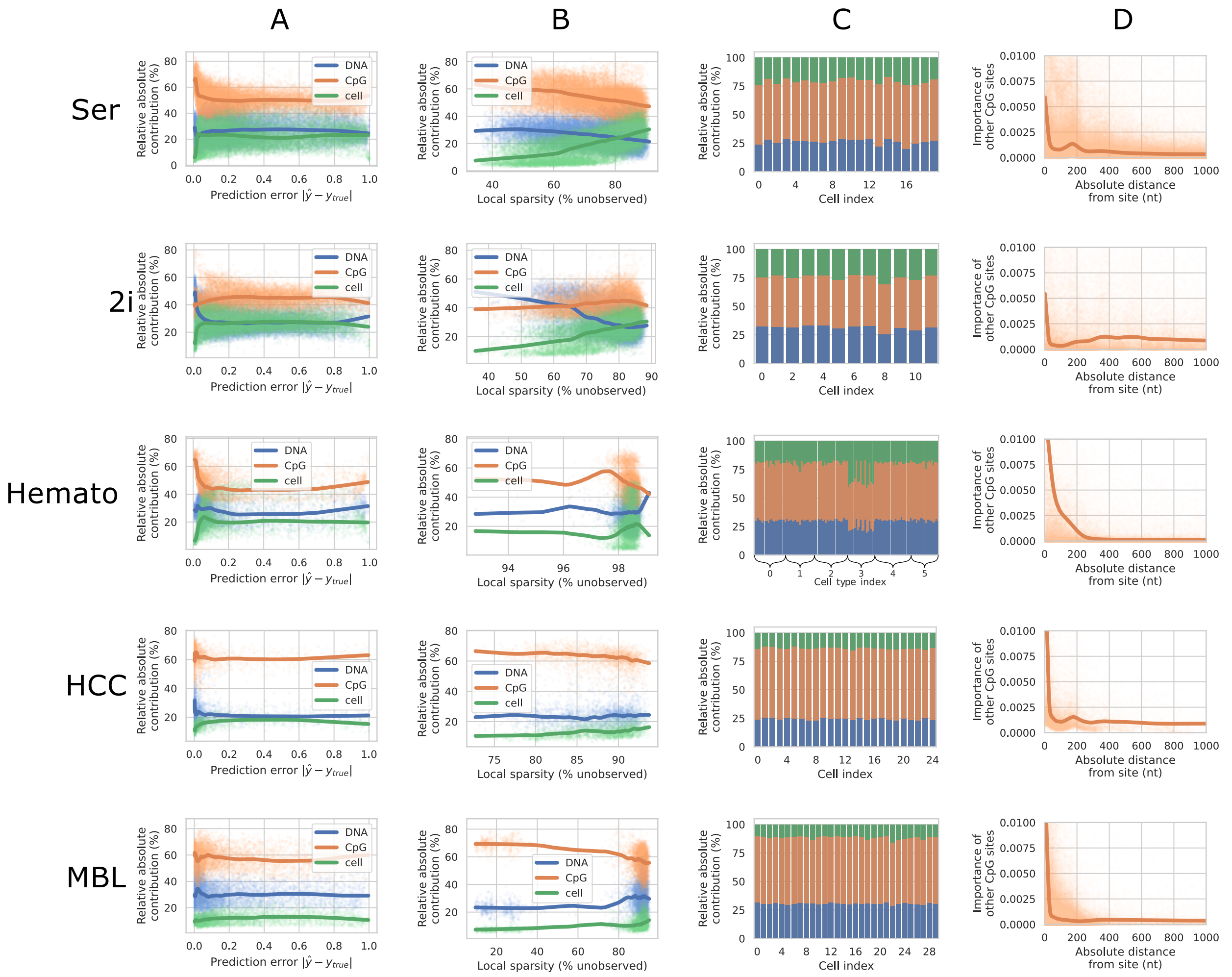


Figure 5: **Integrated gradients interpretation results.** Columns indicate the following: **(A)** Relative absolute contribution of the three input embeddings in function of prediction error. **(B)** Relative absolute contribution in function of local sparsity. **(C)** Relative absolute contribution stratified per cell. **(D)** Contribution of observed CpG sites in function of their absolute distance to the prediction site. Rows indicate the different datasets. From top to bottom: Ser, 2i, Hemato, HCC and MBL. Generally speaking, following statements hold true: (1) CpG contribution is higher when the model is more confident (with a small prediction error), (2) higher sparsity translates in lower contribution of CpG embeddings, (3) differences in relative contributions between cells are negligible, and (4) importance of other CpG sites for prediction decreases with absolute distance from site. A notable exception to statement (3) is found for the Hemato dataset. For this dataset, cell embeddings show a considerably higher contribution for the hematopoietic stem cell type than for the other (progenitor) cell types.
